# Locally distributed abstraction of temporal distance in human parietal cortex

**DOI:** 10.1101/249904

**Authors:** Qun Ye, Yi Hu, Yixuan Ku, Kofi Appiah, Sze Chai Kwok

**Author notes:** Corresponding author (Sze Chai Kwok).

## Abstract

An enduring puzzle in the neuroscience of memory is how the brain parsimoniously situates past events by their order in relation to time. By combining functional MRI, and representational similarity analysis, we reveal a multivoxel representation of time intervals separating pairs of episodic event-moments in the posterior medial memory system, especially when the events were experienced within a similar temporal context. We further show such multivoxel representations to be vulnerable to disruption through targeted repetitive transcranial magnetic stimulation and that perturbation to the mnemonic abstraction alters the neural—behavior relationship across the wider parietal memory network. Our findings establish a mnemonic “pattern-based” code of temporal distances in the human brain, a fundamental neural mechanism for supporting the temporal structure of past events, assigning the precuneus as a locus of flexibly effecting the manipulation of physical time during episodic memory retrieval.

## Introduction

Time in physics is operationally defined as “what a clock reads”. While the passage of time between two moments can be precisely measured by a quartz crystal oscillator or biologically registered by distributed sets of brain regions across intervals of time (1, 2), how the human brain can parsimoniously situate past events by their order in relation to time and *abstract* temporal distances separating events in long-term, episodic memory is incompletely understood (3).

Representations of brief elapsed time can be inferred from single neuron activities in the primate brain (4-6). Time-registering neurons are found to code time with high precision in the cortico-basal ganglia circuits (5), inferior parietal cortex (4) and medial temporal lobe (6) across short timescales. Recent work in rats has provided evidence that temporal information is encoded across time scales from seconds to hours within the overall population state of the lateral entorhinal cortex (7). In contrast, when complex, coherent experiences become consolidated into long-term memories (8), the neural circuits that build time representations as an infrastructure for episodic retrieval are theorized to be distinct from those implicated in hippocampal-dependent encoding (9, 10) and retrieval (11, 12), and from those during transient temporal processing (4–6). For the recollection of long-term autobiographical memories or episodic events, the posterior medial (PM) memory system, including hippocampus, precuneus and angular gyrus, plays an instrumental role (13). Event representations in these regions generalized across modalities (e.g., EEG and MRI) and domains (e.g., perception and memory) (14–16).

Temporal representation is intertwined with the construct of context. A prominent memory model posits that item representations are linked to a changing “context” at encoding, such that a common retrieved context is triggered during recall for items that were experienced within a similar temporal context (17). However, the critical issue of how elapsed time between pairs of long-term episodic events - and its interplay with the encoding context - is represented by the PM system has yet to be addressed. Here we investigated the abstraction, at a macro-anatomical level, of temporal distances that were encoded more than 24 hours previously (18, 19), and determined how several members of this large cortical system are differentially implicated in this putative mnemonic function (20).

Combining functional magnetic resonance imaging (fMRI) with an interactive-Video memory paradigm and a temporal order judgement task (TOJ; **Fig 1A**)—a validated paradigm to study neural correlates underpinning temporal distances between units of memory traces (10, 18, 19)—we adopted a two-forked protocol to ascertain how temporal distances separating pairs of past moments-in-time are represented in the human neocortex. On the one hand, we identified a locally distributed neural representation characterizing the neural patterns of retrieving temporal distances using a multivariate searchlight representational similarity analysis (RSA) (21). We parametrized a large set of pairs of event-moments geometrically separated by varying temporal intervals and applied RSA to compare neural representational dissimilarity matrices (RDM) with a number of parametric, condition-rich hypothetical/candidate models. Applied across the entire brain, the searchlight approach identifies local multi-voxel patterns driven by structured co-activation at a voxel level within the size of the 9-mm radius spherical searchlight, thereby giving us a snapshot of the locally distributed neural architecture supporting temporal order judgements. On the other hand, to enhance the causal strength of the anatomical associations thereby revealed, we focally disrupted the identified critical region with repetitive transcranial magnetic stimulation (rTMS, **Fig 1F**), seeking to confirm its functional necessity for mediating the distributed representation of temporal distances. The spatial scale of rTMS-induced disruption is comparable to that of our chosen searchlight, rendering it an optimal tool for targeted, reversible disruption of the distributed representation of interest.

For memory encoding, participants played an interactive video game containing seven distinct yet related chapters, each in the range of tens of minutes on day 1 (**S1 Fig, S1 Table**). By the nature of the video game, within chapter segments contained more coherent narrative strands than those across chapters, yet all chapters were connected by a common plot. After a 24-hour retention period (day 2), on each trial, participants judged the temporal order of two images (extracted from their individually-played video game, **Fig 1B**), depicting two time-points in their encoded memory, while their blood-oxygen-level-dependent (BOLD) activity was measured (TOJ task, **Fig 1C**). Assuming a scale-free temporal memory representation (22), we manipulated the between-images temporal distances (TD) for all pairs of images so that the TD distribution adhered to a power function permitting scale-invariance across subjects (23) (60 levels of TD, **Fig 1D**). To test the interaction effect between TD and its encoding context, we manipulated the factor “context” by controlling whether the paired images presented at TOJ task were extracted from the same chapters or two adjacent chapters of the video game while keeping the 60 TDs fully matched between the two conditions (Within-chapter vs. Across-chapter, **Fig 1E**).

**Fig 1.**
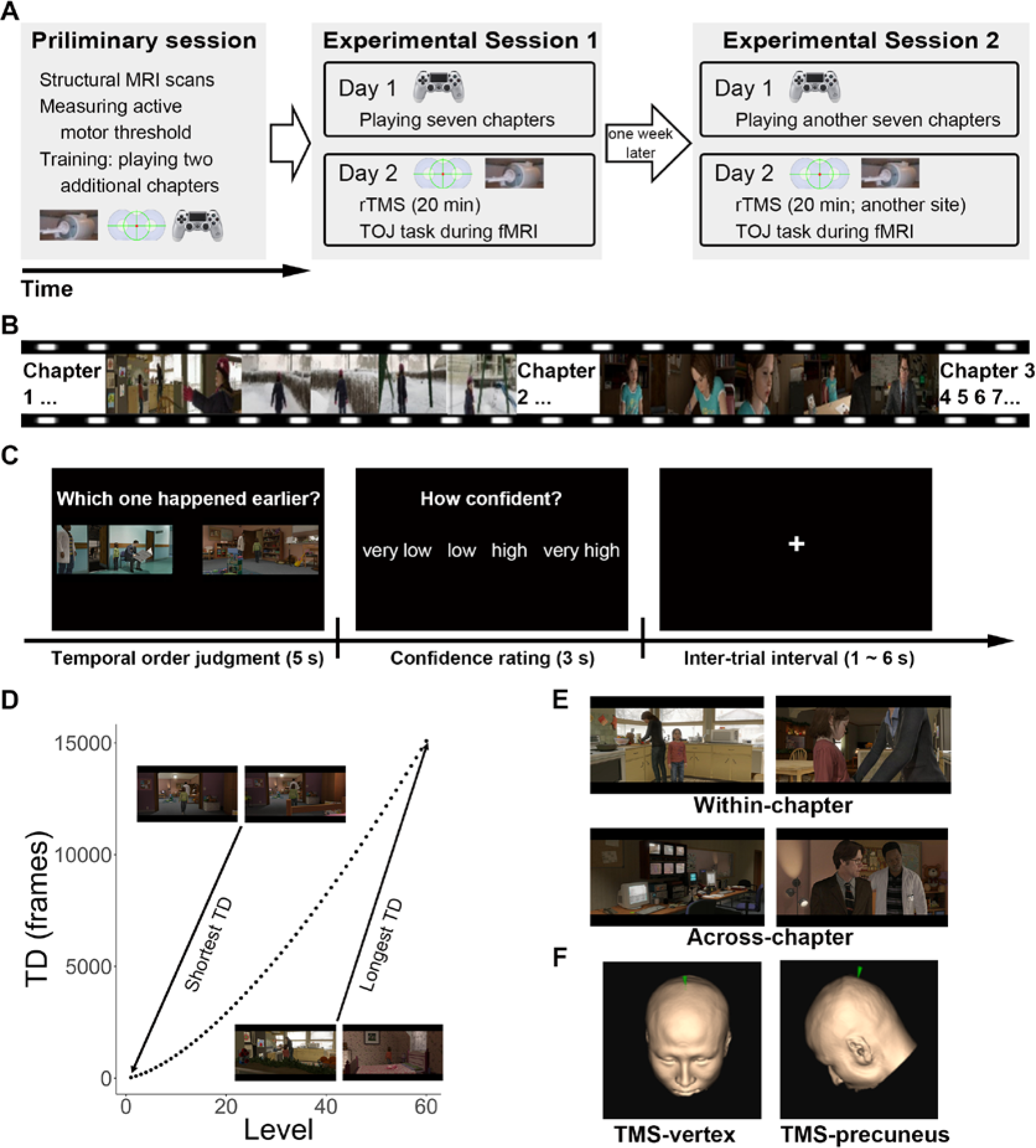
Experiment overview. **(A)** In experimental sessions 1 and 2, participants played a video game containing seven related chapters with a first-person perspective for encoding, and 24 hours later, received 20 min of repetitive transcranial magnetic stimulation (rTMS) to either one of two cortical sites before performing a temporal order judgement task during fMRI. Order of TMS sites (within-subjects) and choices of video game chapters were counterbalanced across subjects (**S1 Table**). The two experimental sessions were conducted on different days to minimize rTMS carry-over effects (mean separation = 8 days). Participants underwent structural MRI scans and familiarized themselves with the gameplay using a console prior to experimental sessions proper. **(B)** Gameplay video: each encoding session consisted of seven chapters (**S1 Fig**). **(C)** Temporal order judgement task. Participants chose the image that happened earlier in the video game and reported their confidence level. **(D)** 60 levels of temporal distances (TD) were generated for each subject according to their subject-specific video-playing duration. Although the absolute TD were different across subjects (**S1 Fig**), we ensured it to be scale-invariant using a power function during image selection. Actual TDs from one subject (subj01) are shown. **(E)** Two pairs of images were extracted from the same chapter (Within-chapter) or two adjacent chapters (Across-chapter). The 60 levels of TD were fully matched within-subjects for these two conditions. Note that scenes depicted in Within-chapter tended to be more contextually similar than those depicted in Across-chapter. **(F)** TMS stimulation sites, superimposed onto one subject’s MRI-reconstructed skull, are marked by a green pointer. The MNI coordinates for precuneus stimulation: x, y, z = 6, −70, 44.

## Results

### Behavioral results

We first looked into the interaction effect between TD and encoding context and the TMS effect on memory retrieval. We collapsed the 60 TD conditions into two levels (short vs. long) for each subject and analyzed the behavioral performance of TOJ (dependent variable: accuracy or reaction times or confidence level) as a function of TMS stimulation site, Context and TD. We ran a three-way repeated-measures ANOVA (TD: Short/Long x Context: Within-chapter/Across-chapter × TMS: TMS-vertex/TMS-precuneus) on task accuracy, and obtained a significant main effect of TD (*F*_(1,16)_ = 25.53, *P* < 0.001, *η*^*2*^ = 10.02%) and a significant two-way interaction effect between TD and Context (*F*_(1, 16)_ = 5.97, *P* = 0.026, *η*^*2*^ = 2.71%). Such interaction effect was driven by a significant difference in accuracy between short and long TD in Within-chapter condition (*t*_(33)_ = 5.94, *P* < 0.001), but not in Across-chapter condition (*t*_(33)_ = 1.61, *P* = 0.117) (**Fig 2A**).

A similar two-way interaction effect was found in reaction times (*F*_(1, 16)_ = 24.21, *P* < 0.001, *η*^*2*^= 1.16%), with longer RT in short than in long TD in Within-chapter condition (*t*_(33)_ = −3.33, *P* = 0.002) but longer RT in long TD condition in Across-chapter condition (*t*_(33)_ = 2.83, *P* = 0.008) (**Fig 2B**). No three-way interaction effects were found in either of the measures (*P*s > 0.05). A similar behavioral pattern was found in the measure of confidence level (**S2 Fig**). In terms of TMS effect on behavioral measures, TMS to the precuneus resulted in slowed reaction times (main effect of TMS: *F*_(1, 16)_ = 5.34, *P* = 0.035, *η*^*2*^= 2.65%) as compared with the TMS-vertex condition.

**Fig 2.**
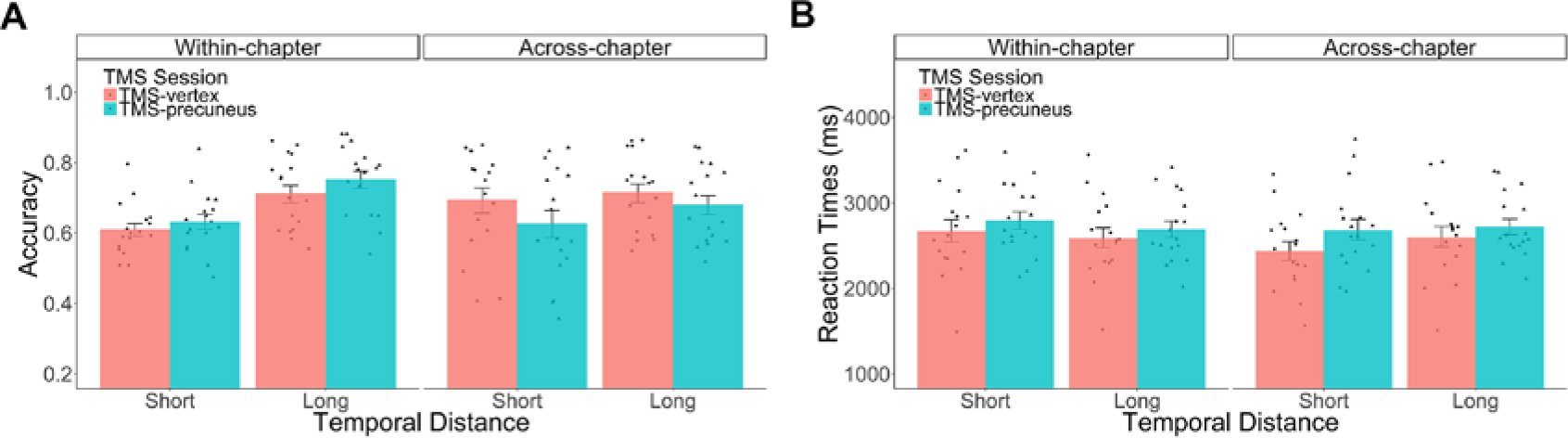
Behavioral results. **(A)** Accuracy: The difference between short and long temporal distances was more pronounced for Within-chapter condition than for Across-chapter condition irrespective of TMS stimulation. **(B)** Reaction times: The differences between short and long temporal distances were in an opposite way between the Within-chapter condition and the Across-chapter condition. TMS stimulation resulted in longer reaction times in TOJ task. Dots denote behavioral performance per subject. Error bars denote the SEM over subjects.

### Functional MRI results

#### Representation of temporal distances

First and foremost, we searched for neural representations which might resemble the matrix of temporal distances using searchlight representational similarity analysis (21). Without *a priori* bias for any region of interest, we searched the entirety of the cortex using an RDM consisting of 60 levels of logarithmically-transformed subject-specific TD (**Fig 3A**) and identified clusters of voxels that contain information of the set of geometrically defined temporal distances in memory (see **Materials and methods** and **S3 Fig**). Within this 60 × 60 RDM of temporal-distances, we revealed that the neural pattern of judging the temporal order of a pair of memories separated with a given temporal distance is more similar to other temporal order judgements which also enclosed temporal distances of a comparable scale. These voxels were in the posteromedial parietal areas, bilateral angular gyri, and middle frontal gyri (**Fig 3B**, **S2 Table**).

The temporal-distance memory representation could be confounded by perceptual similarity in each pair of images. To address this concern, we conducted six separate RSAs, in which we indexed perceptual similarity between the image-pairs by six different metrics, some drawn from the visual categorization models referred by Greene et al (24) and Aminnoff et al (25). The six RDMs are Red Green and Blue (RGB) cross-correlation RDM, RGB-intensity RDM, RGB-histogram RDM, Scale Invariant Feature Transform (SIFT) RDM (26), Speeded Up Robust Features (SURF) RDM (27), and Histogram of Oriented Gradients (HOG) RDM (28) (see **Materials and methods**). No similar representation was observed in the posteromedial parietal cortex using these candidate RDMs except the SURF RDM (**S4B-G Fig**). Importantly, to further exclude the contribution of these additional perceptual properties, we conducted a whole brain searchlight correlation between the TD RDM and the brain response pattern, this time regressing out the six perceptual RDMs using a partial Spearman correlation analysis method. This partial correlation analysis showed that the effects of abstraction of TD distributed in the posteromedial parietal areas remained even after removing the influences of all of these perceptual properties, suggesting that TD multivoxel representation could not be driven simply by the perceptual properties (**S4J Fig**).

Considering previous work on space-time relationships in episodic memory (11, 29), we also quantified the space displacement embedded between the image-pairs by computing the number of locations each participant had virtually traversed in the video game, and entered them into a subject-specific Situational Changes RDM (**S5 Fig**). Despite the space-time correlation in the encoding material (**S8 Fig**), subject-specific Situational Changes RDMs explained very little variances in comparison to our temporal-distance RDM (**S4A Fig**). We also confirmed that participants’ behavioral performance RDMs, namely reaction times and accuracy, could not explain the putative temporal-distance representation (**S4H-I Fig**).

Having identified a multivariate pattern underlying the temporal memory abstraction in the posterior medial parietal cortex, we asked further whether there were voxels whose activities change monotonically as a function of temporal distance using a standard univariate approach, irrespective of its multivoxel characteristics. A whole-brain parametric modulation analysis revealed TD-specific BOLD signals in a cluster within the posteromedial region, including the precuneus (**Fig 3C**, **S3 Table**). This parametric relationship could not be attributed to difficulty (results were the same after trial-by-trial reaction times were regressed out, **S6A Fig**). Importantly, since the two types of analyses extracted two different kinds of neural information, their overlap in the precuneus jointly confirms the critical involvement of this region. We accordingly created a conjunction map (**Fig 3D**), so that both the multivariate and univariate results underlying the temporal distance abstraction would be available for the next analysis.

We then performed inference tests to statistically assess whether the TD RDM is significantly correlated with subject-specific neural RDM (labelled as reference RDM in **Fig 3**) estimated from the fMRI response pattern in accordance to the 60 TD levels (4 repetitions per TD level) within this conjunction region. This would allow us to test whether the TD RDM could explain the neural representation better than the other candidate RDMs. The TD RDM (our best-performing model) accounted for the variance far better than the other candidate RDMs, such as SC RDM, six perceptual RDMs and two behavioral RDMs (**Fig 3E-F**). These results showed that the neural signals coded in the posteromedial region as revealed by these searchlight analyses were most attributable to the mnemonic representation of temporal distances.

**Fig 3.**
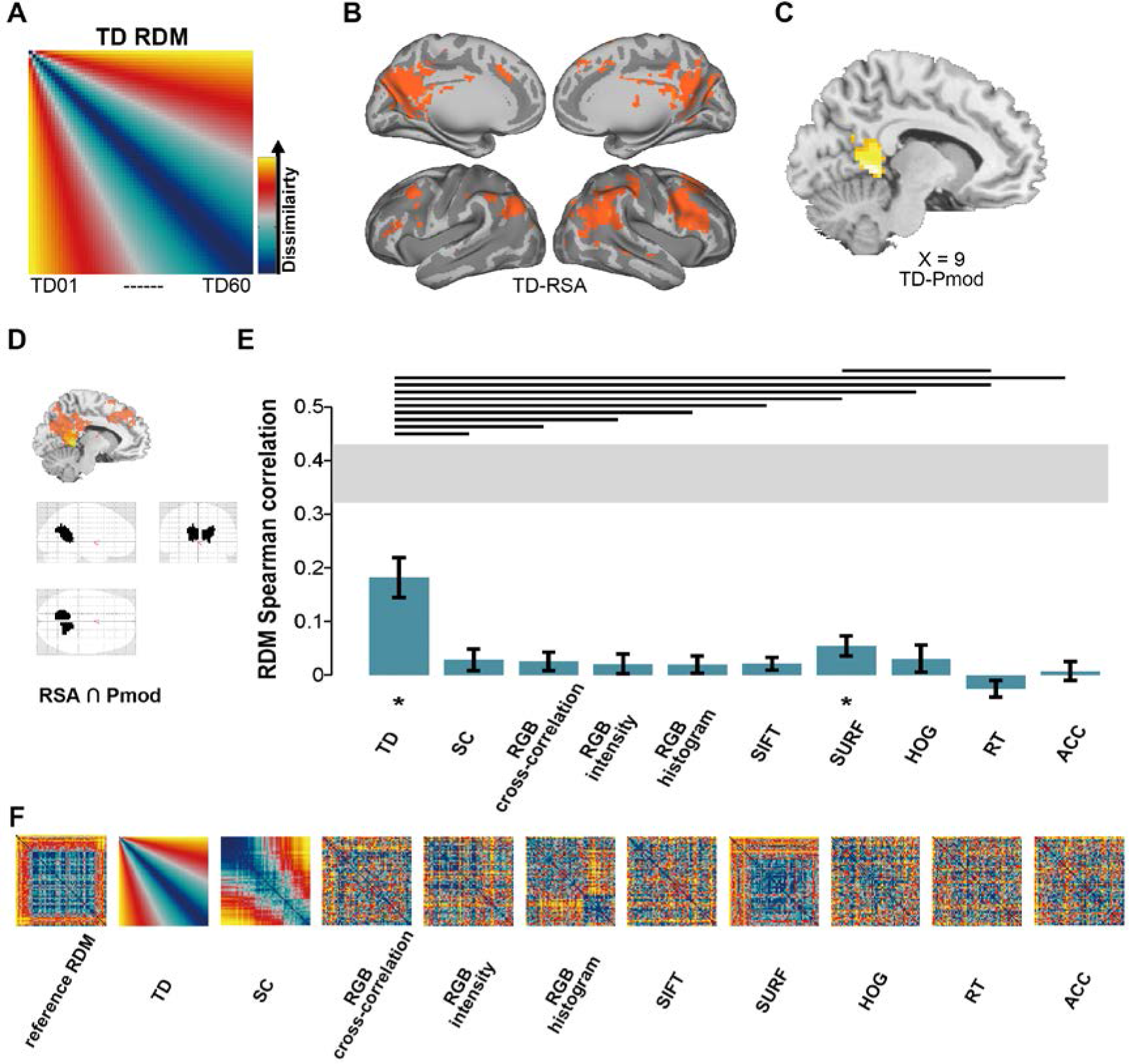
Representation of temporal distances. Abstraction of temporal distances in posteromedial parietal cortex. **(A)** TD representational dissimilarity matrix (RDM) for searchlight RSA. The RDM consisted of 60 subject-specific TD levels. Any two event-moments that are separated by short TD will get increasingly dissimilar with other two event-moments as the TD increases. **(B)** Using TD RDM for searchlight RSA, clusters of voxels that contained TD information were primarily in the posteromedial cortex, bilateral angular gyri, and bilateral middle frontal gyri. **(C)** Activation signal intensity from parametric modulation analysis (Pmod). The intensity of these voxels, primarily in the left precuneus, increased as a function of TD. **(D)** Conjunction region for neural signals extraction for subsequent inferential tests. The mask was created by intersecting the similarity map from RSA and activation map from Pmod (**S2&3 Table**). **(E)** Inferential comparisons of multiple model representations. Several candidate RDMs were tested and compared for their ability to explain the neural reference RDM extracted from the conjunction mask. TD RDM was the best model among these candidate RDMs to explain the variances in the neural reference RDM. Horizontal lines over two bars indicate that the two models having significant statistical differences (signed-rank test) after multiple testing correction. The gray horizontal bar indicates the expected performance of an unknown true model, given the noise in the data. Asterisks indicate a significant Spearman correlation between the subject-specific reference RDM and candidate RDMs. Error bars indicate the SEM over participants. * *P* < 0.05, FDR-corrected. **(F)** Reference (neural) RDM and candidate RDMs. The reference RDM was estimated using the BOLD responses elicited by the stimuli during fMRI in accordance to the 60 TD levels in the conjunction region. We averaged 17 subject-specific neural reference RDMs for display purpose. SC RDM was computed by the number of locations each participant had virtually traversed in the video (**S5 Fig**). Perceptual RDMs (RGB cross-correlation, RGB intensity, RGB histogram, SIFT, SURF,HOG) were computed based on image properties (see details in Materials and methods). Behavioral RDMs (RT and ACC) were computed based on the behavioral performance of reaction times and percentage correct for each trial (see details in Materials and methods). MRI results are displayed at *P*_uncorrected_ < 0.001.

#### TD representation is context-dependent

To test the hypothesis that the TD-neural pattern similarity index to be higher when the two images are extracted from a “similar context” than when they are from two “different contexts”, we ran a new searchlight RSA, now separately for the Within-chapter and Across-chapter trials. The representation of TD was observed only in the Within-chapter condition (**Fig 4A**, left) but not in the Across-chapter condition (**Fig 4A**, right). The voxels identified by the searchlight RSA were in the precuneus, retrosplenial cortex, and angular gyri bilaterally.

For statistical inference, we extracted the similarity index within the aforementioned conjunction mask (RSA and pmod maps), using a bias-free leave-one-subject-out method (see **Materials and methods**). In line with our prediction, the voxels in the Within-chapter trials contained higher pattern similarity to the TD RDM than Across-chapter trials (**Fig 4A**, middle panel; one-tailed: *P* = 0.04), confirming the neural pattern similarity related to the TD RDM was indeed stronger in Within-chapter trials. We then performed statistical inference tests to assess whether the TD RDM explains the neural reference RDM better than other candidate RDMs separately for Within-chapter and Across-chapter conditions (**Fig 4B**). The TD model accounted for the variances far better than the other candidate RDMs in the Within-chapter condition (**Fig 4B**, left). By contrast, the TD RDM (and all other candidate RDMs) failed to explain the neural reference RDM in the Across-chapter condition (**Fig 4B**, right).

This context-dependent difference was also found in a voxel-wise univariate analysis. The beta-estimates (*β*) from a pmod analysis using TD as a regressor were significantly higher in the Within-chapter condition compared to the Across-chapter condition (**Fig 4C**). These results were consistent in a control analysis while RT were regressed out from the pmod analysis (**S6B Fig**). This confirmed that the mnemonic representation of temporal distances was determined by whether the pairs of images were experienced within a similar context, corroborating the interaction between temporally- and semantically-defined factors observed during memory encoding (9) and retrieval (12).

**Fig 4.**
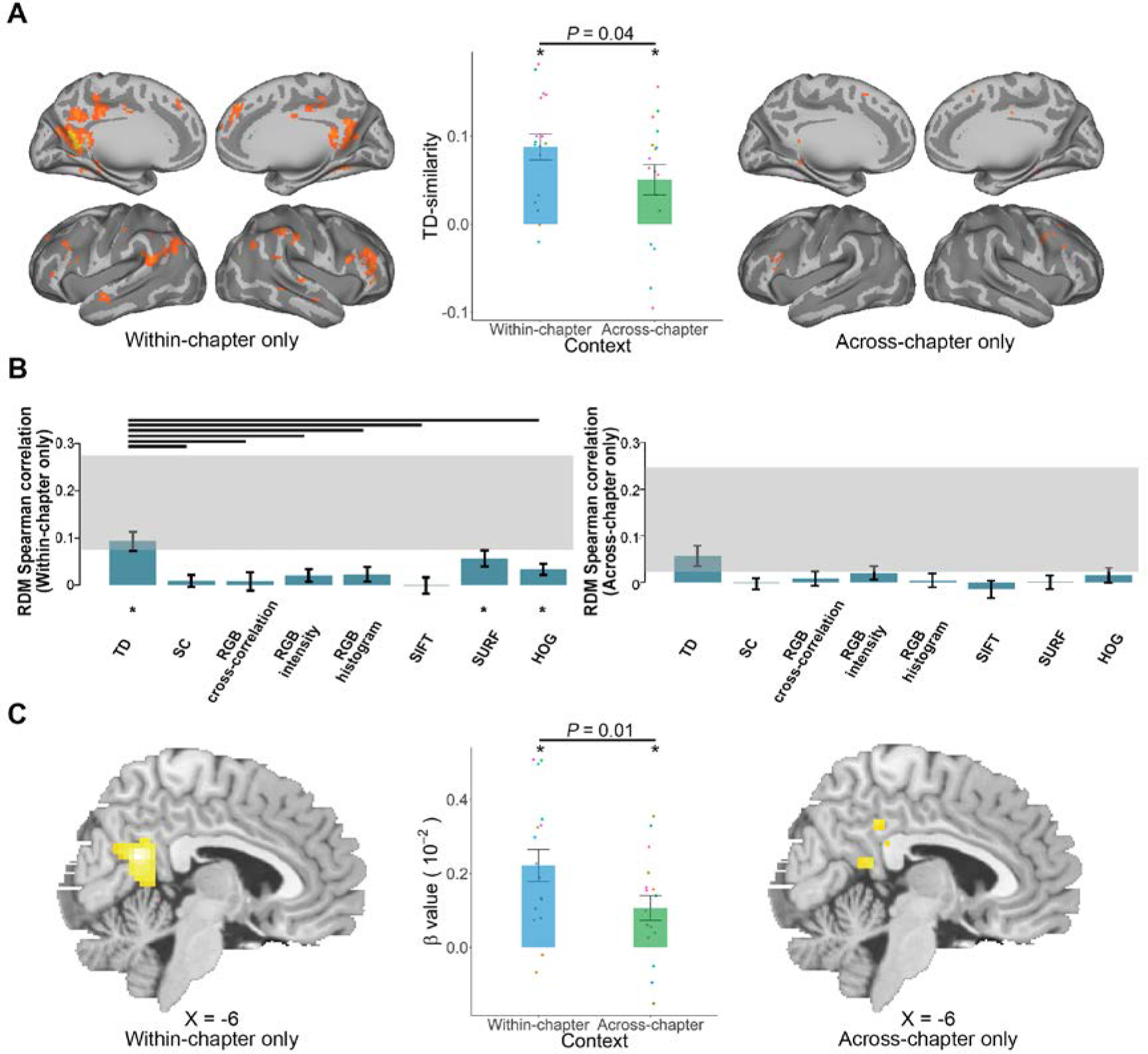
TD representation is context-dependent. **(A)** Stronger multivoxel similarity representing TD variation in the Within-chapter condition than in the Across-chapter condition in the conjunction region (middle panel, one-tailed: *P* = 0.04). Each same color dot represents one subject. Error bars denote the SEM over subjects. * *P* < 0.05, FDR-corrected, one-sample t-test against zero. **(B)** Separate model comparisons showed the TD RDM is the best model to explain neural variances in the Within-chapter condition (although SURF RDM and HOG RDM are also related to the neural reference RDM), whereas all the candidate RDMs could not explain the neural variances in the Across-chapter condition. **(C)** Stronger parametric activation intensity in the Within-chapter condition than in the Across-chapter condition.

#### TMS reduced the precuneal representation of TD

To causally ascertain a pivotal role of the precuneus in this memory operation we deployed a disruptive technique, strategically targeting the precuneus with repetitive transcranial magnetic stimulation to interrogate changes on both neural and behavioral levels (within-subjects: TMS-precuneus vs. TMS-vertex; **Fig 1F** and **S1 Table**). We found the widespread representation of TD disappeared following TMS on the precuneus, either considering the Within-chapter condition (**Fig 5A**) or collapsing Within-chapter and Across-chapter conditions. Model comparison results showed that the correlation values between all the candidate RDMs and neural reference RDM now failed to reach statistical significance even when only considering Within-chapter trials alone (**Fig 5B**). In contrast, the activation-based Pmod analyses showed that TMS to the precuneus did not induce any discernable changes in the univariate BOLD intensity (**S6C Fig**)

In light of the fractionation view for the parietal cortex (30), we further tested the possibility that there might be differences in the patterns of neural activity associated with the abstraction of temporal distances in the sub-regions of the PM memory network (13). Based on our main MRI results (**Fig 3B**) and previous work on the parcellation of the PM memory network (20), we have chosen six anatomical regions-of-interest in the PM memory network (ROIs: bilateral precuneus, bilateral angular gyrus and bilateral hippocampus; see **Materials and methods**), together with the primary visual region (entire occipital cortex) as a control, to more finely characterize the disruptive effect caused by the TMS.

We extracted the similarity indices from these ROIs and found that the neural-TD pattern similarity in the left precuneus was significantly weakened following TMS to the precuneus specifically for Within-chapter condition (**Fig 5C**, one tailed: *P* = 0.02; to a lesser extent, also the right precuneus and left angular gyrus, **S7A&D Fig**). Specifically, we found that changes in individuals’ neural-TD pattern similarity in the vertex condition to be associated positively with their TOJ memory performance in this key region (**Fig 5C**, *r* = 0.60, *P* = 0.04; also in the left hippocampi, see **S7B Fig**), implying these multivoxel representations are relevant neurobiological prerequisites for the ability to support temporal order judgement. In the experimental condition wherein we disrupted the precuneal activity with magnetic field prior to retrieval, changes in neural-behavioral correlation were resulted (**Fig 5C**, right panel; comparison between two correlations: *z* = 2.07, *P* = 0.04). Since the focal perturbation altered the mnemonic representation across parts of the PM system implicating the precuneus, the angular gyri and the hippocampi, the putative disruption might have been effective through inducing alternation in functional connectivity between multiple regions, or more globally throughout the entire parietal memory network (31, 32).

**Fig 5.**
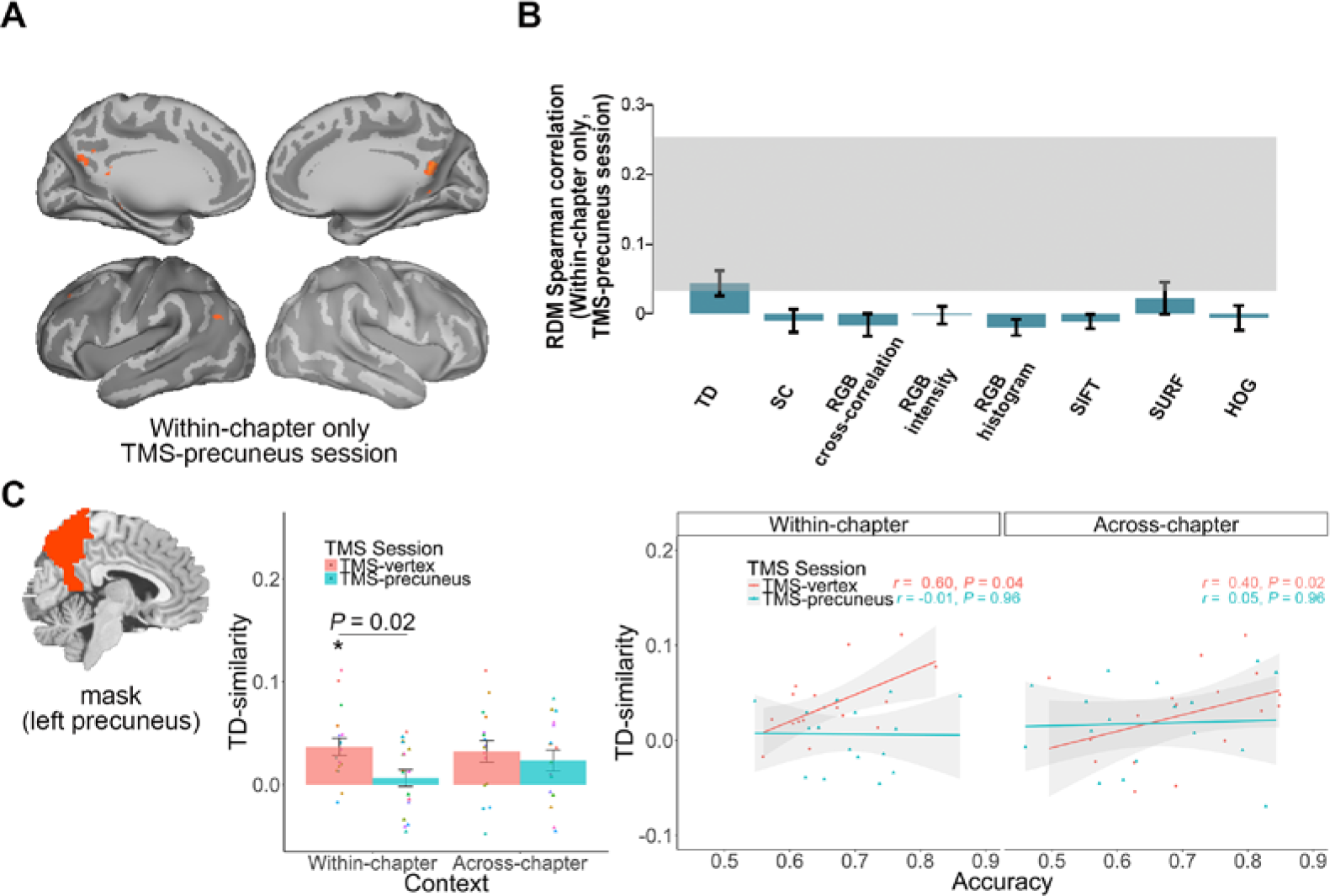
TMS reduced the precuneal representation of TD. TMS reduced the precuneal representation of TD. **(A)** Very few voxels survived in the searchlight RSA after TMS on the precuneus (voxel level: *P*_uncorrected_ < 0.001; cluster level: *P*_FWE_ < 0.05). **(B)** Model comparisons showed that the candidate RDMs explain little neural variances after TMS on the precuneus. **(C)** We used left precuneus as a mask to extract TD-similarity values from subjects and found that TMS-precuneus, compared to the TMS-vertex, reduced the TD representation specifically in the Within-chapter condition (middle panel, one tailed: *P* = 0.02). Further correlation analyses showed that TMS on the precuneus weakened the significant neural-behavioral correlation (TMS-vertex session: *r* = 0.60, *P* = 0.04; TMS-precuneus session: *r* = −0.01, *P* =0. 96; comparison between two correlations: *z* = 2.07, *P* = 0.04). *P* values are FDR-corrected.

## Discussion

By combining fMRI, rTMS and multivoxel pattern analysis with a novel interactive memory paradigm, we provided the first characterization of multivoxel pattern of temporal distance between pairs of episodic events in the human parietal cortex and showed that BOLD intensity of posteromedial cortex, especially the precuneus, varied as a function of temporal distance. We showed that such information carried by these regions is more pronounced within a similar context and such representations reduced significantly after the precuneus was perturbed. We further revealed that the precuneal representation of TD is associated with subjects’ memory performance, especially when two images for temporal order judgment were extracted from the same context.

Our findings align with the Temporal Context Model (17) which stipulates that the fine-grained TD memory information distributed in this cortex is stronger when paired images were associated within a similar context. The parietal representation of temporal distances between pairs of episodic events observed here, as revealed by both univariate and multivariate pattern analyses, might act in parallel with hippocampal cells that code specific moments in time or temporal positions (33), or act independently as a separate mnemonic establishment of episodes over and above the hippocampal memory ensemble (34).

Previous studies have associated neural similarity in the hippocampus with both spatial and temporal aspects of episodic memory (29, 35, 36). Studies on rats have also showed that gradually changing representation, as manipulated by temporal lag/distance, is manifested in the hippocampus (10). These studies have mostly focused on neural similarities in the hippocampus (29, 36, 37), whereas here we revealed a TD representation primarily in the PM system. This discrepancy might be due to the fact that instead of using a passive paradigm (viewing movies or listening to narratives), we used an active paradigm (first-person perspective gameplay for encoding), which should have implicated a distinct system upon retrieval (38),(39) when the memories in question are akin to real-life autobiographical/episodic experiences. The consideration that a temporal order judgment task was used in which participants had to extract temporal distance for making a decision (40), and then the neural signals were assessed in relation to the actual temporal distances rather than to subjective estimation of temporal separation might have also engaged the PM memory system more heavily.

It is theoretically interesting that the multivariate representations were more vulnerable to the magnetic stimulation than the voxel-wise signal intensity (note that TMS to the precuneus did not induce any discernable changes in univariate BOLD level, see **S6C Fig**). These findings support our argument that memory traces that are represented during temporal order judgement are indeed conveyed in some localized multivoxel readouts housed in the PM system cortices, above and beyond the modulated changes in canonical BOLD activation. Building on extant connectivity findings between the hippocampus and neocortical regions (13, 41, 42) and the hippocampal role in temporal context memory (9, 12), our demonstration of distributed pattern of temporal information in the posteromedial parietal region implied the existence of a higher level parietal mnemonic readout of temporal distances between episodic experiences.

Model comparisons incorporating several control analyses robustly confirm that such precuneal representation of temporal distances do not merely reflect some trivial effects related to task difficulty (18, 43) or inferred distance based on image properties. Although some perceptual models (such as SUFT model) might produce a similar RSA map, when we partially removed the respective influences of perceptual properties, the effects of TD representation in the posteromedial parietal cortex were largely unaffected (**S4J Fig**). Moreover, in terms of the stimuli features, since our images were extracted randomly across chapters of the video game (**S1C Fig**), and since the TD distributions were fully matched between the Within-chapter and Across-chapter conditions, predictions made by the positional coding theory (44) would also not be sufficient to account for the differential TD representational results across these two conditions.

In summary, our multivariate searchlight results reveal that the temporal distance representations in the posterior parietal cortex, especially in the precuneus, during TOJ retrieval are determined by how temporally distant (and how similar the encoding contexts) two given event-moments the subjects had encountered (18, 19). We also establish that this multivoxel mnemonic abstraction is localized in the precuneal area and perturbation to it alters the neural—behavior relationship across the global parietal memory network, assigning this structure as a locus of flexibly effecting the manipulation of physical time during episodic memory retrieval.

## Materials and methods

### Participants

Twenty individuals participated in the study (7 female, 22.55 ± 1.54 years, mean ± sd). Data from 3 subjects were excluded due to either poor performance (1 subject performed at chance level) or scanner malfunction (projector crashed during scanning for 2 subjects at TMS-vertex session), resulting in a final group of 17 subjects (7 female, 20.65 ± 1.54 years, mean ± sd). All subjects were unfamiliar with the video game, had normal or correct-to-normal vision and did not report neurological or psychiatric disorders or current use of psychoactive drugs. All subjects were eligible for MRI and TMS procedures based on standard MRI safety screening as well as on their answers to a TMS safety-screening questionnaire (45). No subjects withdrew due to complication from the TMS or MRI procedures, and no negative treatment responses were observed. All subjects gave written informed consent and were compensated for their participation. All procedures were performed in accordance with the 1964 Helsinki declaration and its later amendments and approved by University Committee on Human Research Protection of East China Normal University (UCHRP-ECNU). The number of participants was determined based on previous studies with similar design (9, 32).

### Experimental design, stimuli, and tasks

#### Encoding: Interactive video game

The action-adventure video game (Beyond: Two Souls) was created by the French game developer Quantic Dream and played in the PlayStation 4 video game console developed by Sony Computer Entertainment. Participants played the game using a first-person perspective. To ensure that the participants mastered the operational capability, they were trained to play the game with two additional game chapters (Training chapters: Welcome to the CIA, and The Embassy). The training session varied in duration depending on the dexterity of each participant on using the console (40 - 60 min per chapter). After the training session, participants played 14 chapters in total across two sessions: 7 in Experimental Session 1 and then another 7 in Session 2 (**Fig 1**). The video game they played were recorded and stored as a single video file in MP4 format (Chapters 1~7: My Imaginary Friend, First Interview, First Night, Alone, The Experiment, Night Session, Hauntings; Chapters 8~14: The Party, Like Other Girls, Separation, Old Friends, Norah, Agreement, Briefing; see **S1 Fig**).

#### Retrieval (scanned): Temporal Order Judgment (TOJ) task

The TOJ retrieval task required participants to choose the image that happened earlier in the video game they had encoded. The task was administrated inside an MRI scanner, where visual stimuli were presented using E-prime software (Psychology Software Tools, Inc., Pittsburgh, PA), as back-projected via a mirror system to the participant. Each trial was presented for 5 s during which participants performed the temporal order judgment. They were then allowed 3 s to report their confidence level following the memory judgement. Participants performed the TOJ task using their index and middle fingers of one of their hands via an MRI compatible five-button response keyboard (Sinorad, Shenzhen, China). Participants reported their confidence level (“Very Low”, “Low”, “High”, or “Very High”) regarding their own judgment of the correctness of TOJ with four fingers (thumb was not used) of the other hand. The left/right hand response contingency was counterbalanced across participants. Participants were told they should report their confidence level in a relative way and make use of the whole confidence scale. Following these judgments, a fixation cross with a variable duration (1 - 6 s) was presented. Each participant completed 240 trials in each of the two experimental sessions. Participants were given 15 practice trials using paired images extracted from the two additional chapters they had played in the training session out of the scanner to ensure they understand the task procedure. Participants completed a surprise recognition test after TOJ task outside scanner; data of which are not reported here.

For the TOJ task, we selected still images from the subject-specific recorded videos which the participants had played the day before. Each second in the video consisted of 29.97 static images (frames). For each game-playing session, 240 pairs of images were extracted from the seven chapters and were paired up for the task based on the following criteria: (1) the two images had to be extracted from either the same chapters or adjacent chapters (Within-chapter vs. Across-chapter); (2) the temporal distance (TD) between the two images were matched between Within- and Across-chapter condition; (3) in order to maximize the TD, we first selected the second longest chapter of the video and determined the longest TD according to a power function (power = 1.5), at the same time ensuring the shortest TD to be longer than 30 frames. We generated 60 progressive levels of TD among these pairs (each level repeated twice). In sum, three within-subjects factors regarding the TOJ retrieval task were manipulated: (1) 60 TD levels permitting scale-invariance across subjects between two images (see below); (2) Context (two images extracted from either Within- or Across-chapter); (3) TMS stimulation (TMS-precuneus vs. TMS-vertex, see below).

#### Selection of 60 levels of temporal distances (TDs)

In order to maximize the range of all TDs, we first selected the second longest chapter of the video game and determined the longest TD (*L*), while ensuring the shortest TD to be longer than 30 frames. The 60 TD levels were selected according to this function,

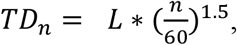

where *L* denotes duration of the second longest chapter of the video game in each experimental session, *n* denotes TD level, and value of *TD*_*n*_ were rounded to the nearest integer using the “round” function in MATLAB. Note that the actual TDs were different across subjects, but since we applied a power function, the scale was thus rendered invariant (22). Image-pairs extraction from each of the chapters were independently conducted across subjects. The numbers of images-pairs extracted from each of the chapters were approximately equal within-subjects.

### Transcranial magnetic stimulation

#### TMS procedure and protocol

TMS were applied using a 70 mm Double Air Film Coil connected to a Magstim Rapid2 (The Magstim Company, Ltd., Whitland, UK). In order to localize the target brain regions precisely, we obtained individual anatomical T1-weighted magnetic resonance images and then imported them into BrainSight (Rogue Research Inc., Montreal, Canada) for stereotaxic registration of the TMS coil with the participants’ brain. The position of the coil and the subject’s head were co-registered with BrainSight, and monitored using a Polaris Optical Tracking System (Northern Digital, Waterloo, Canada) during TMS. Positional data for both rigid bodies were registered in real time to a common frame of reference and were superimposed onto the reconstructed three-dimensional MRI images of the subject using the BrainSight. The center of the coil was continuously monitored to be directly over the site of interest. For all sites (vertex, precuneus, and motor areas for measuring active motor threshold), the TMS coil was held tangential to the surface of the skull and was placed in a rostro-caudal direction. An adjustable frame was used to hold the TMS coil firmly in place, while the participants rested their heads on the chin rest. Head movements were monitored constantly by BrainSight and were negligible. We measured subjects’ active motor threshold, defined as the lowest TMS intensity delivered over the motor cortex necessary to elicit visible twitches of the right index finger in at least 5 out of 10 consecutive pulses. The location used to determine the active motor threshold was identified with a single pulse of TMS over the motor cortex at the left hemisphere. The TMS coil was systematically moved until the optimal cortical site was located to induce the largest and most reliable motor response; this stimulus output was then recorded. The TMS intensity was then calibrated at 110% of individual active motor threshold (stimulator output: 75.2 ± 6.9%, mean ± se, range from 63% to 88%, **S1 Table**). In Experimental Session 1 and 2, the TMS was applied at a low-frequency rate of 1 Hz with an uninterrupted duration of 20 min.

#### TMS stimulation sites

The target stimulation was delivered to the precuneus (18) (MNI x, y, z = 6, −70, 44), whereas the control stimulation was delivered to the vertex. The vertex was defined individually by the point of the same distance to the left and the right pre-auricular, and of the same distance to the nasion and the inion. Due to the folding of the two cerebral hemispheres, the stimulated vertex site lies at a considerable distance from the TMS coil, thereby diminishing the effectiveness of the magnetic pulses. Stimulating the vertex is not known to produce any memory task-relevant effects and deemed as a reliable control site. Stimulation magnitude and protocols in the present study were comparable to those used in similar studies that are robust to produce significant memory-related changes by targeting at the precuneus (46–48) or lateral parietal cortices (31, 32). Immediately after the end of the stimulation, participants performed four runs of Temporal Order Judgment task in the MRI scanner (delay period between the end of TMS and the beginning of MRI: *M*_precuneus_ = 15.29 min, *M*_vertex_ = 20.76 min, *t* (16) = −0.87, *P* = 0.4).

### MRI data acquisition and preprocessing

#### Data acquisition

All the participants were scanned in a 3-Tesla Siemens Trio magnetic resonance imaging scanner using a 32-channel head coil (Siemens Medical Solutions, Erlangen, Germany) at ECNU. In each of the two experimental sessions, a total of 1,350 fMRI volumes were acquired for each subject across 4 runs. The functional images were acquired with the following sequence: TR = 2000 ms, TE = 30 ms, field of view (FOV) = 230 × 230 mm, flip angle = 70°, voxel size = 3.6 × 3.6 × 4 mm, 33 slices, scan orientation parallel to AC-PC plane. High-resolution T1-weighted MPRAGE anatomy images were also acquired (TR = 2530 ms, TE = 2.34 ms, TI = 1100 ms, flip angle = 7°, FOV = 256 x 256 mm, 192 sagittal slices, 0.9 mm thickness, voxel size = 1 × 1 × 1 mm).

#### Preprocessing

Preprocessing was conducted using SPM12 (http://www.fil.ion.ucl.ac.uk/spm). Scans were realigned to the middle EPI image. The structural image was co-registered to the mean functional image, and the parameters from the segmentation of the structural image were used to normalize the functional images that were resampled to 3 × 3 × 3 mm. The realigned normalized images were then smoothed with a Gaussian kernel of 8-mm full-width half maximum (FWHM) to conform to the assumptions of random field theory and improve sensitivity for group analyses (49). Data were analyzed using general linear models and representational similarity analyses as described below with a high-pass filter cutoff of 256 s and autoregressive AR(1) model correction for auto-correlation.

### Functional MRI data analysis

#### Parametric modulation analysis

First-level models were performed on the fMRI data collected from the TMS-vertex session only (either all the trials altogether or separately for Across-chapter vs. Within-chapter conditions). In all of these models, each of the 240 trials was modeled with a canonical hemodynamic response function as an event-related response with a duration of 5 s.

For the TMS-vertex session as a whole (Across-chapter and Within-chapter trials collapsed), we performed two parametric modulation analyses (pmod), each with a different combination of modulatory regressor/regressors (namely, TD; TD + RT). For the TD pmod, we assigned the actual TD values at encoding as the modulatory parameter, and used the polynomial function up to first order. Several regressors of no interest were also included: 6 head movement regressors and 1 missing trial regressor (i.e., no-response trials; number of missing trials of Across-chapter condition: 5.65 ±6.96, of Within-chapter condition: 5.29 ± 6.8; n = 17, mean ± sd) and the run mean. The purpose of this analysis was to test for any linear TD-dependent modulation of signal intensity in the brain between the TD between the two images at encoding and the brain activity during TOJ retrieval of the same events. For the TD + RT pmod, we aimed to identify the voxels whose activities changed as a function of TD after the removal of the influence of reaction times. Each subjects’ RTs corresponding to each TD level were entered as the modulatory parameter, together with the regressors of no interest as above.

For the Across-chapter vs. Within-chapter comparison, we also performed two pmod analyses with identical sets of regressors as described above (namely, TD; TD + RT). We looked for changes in brain responses as a linear function of the regressor of interest (i.e., TD). Maps were created by multiple regression analyses between the observed signals and regressors. The contrast maps from the first-level model of parametric analyses were taken for second-level group analyses and entered into one-sample *t*-tests. The group analyses were performed for each contrast using a random effects model(50). The statistical threshold was set at *p* < 0.05 (FWE corrected) at cluster level and *p* < 0.001 at an uncorrected peak level according to the SPM12 standard procedure. The activation cluster locations were indicated by the peak voxels on the normalized structural images and labeled using the nomenclature of Talairach and Tournoux (1988) (51).

#### Searchlight Representational Similarity Analysis (searchlight RSA)

RSA were conducted using the RSA toolbox (http://www.mrc-cbu.cam.ac.uk/methods-and-resources/toolboxes/) on the fMRI data following realignment and normalization, but without smoothing. In the Across-chapter vs. Within-chapter comparison, each unique TD level was modeled with a separate regressor and was contrasted to produce a T-statistic map (spmT maps), creating 120 statistical maps in total (Across- vs. Within-chapter conditions; 2 repetitions for each TD level). For the TMS-vertex vs. TMS-precuneus comparison, we collapsed the respective trials within the Across- and Within-chapter conditions and generated 60 statistical maps in either of the two sessions (4 repetitions for each TD level). Using searchlight RSA, spherical searchlights with a radius of 9 mm (93 voxels, volume = 2,511 mm^3^) were extracted from the brain volume and then the data (i.e., signal intensity) for the 60 TD levels were Person product-moment (1 - *r*) correlated with every other level to generate a representational dissimilarity matrix (RDM), reflecting the between-condition dissimilarity of BOLD signal response. These neural RDMs were then Spearman-rank correlated with a set of candidate RDMs (see **Fig 3F**), reflecting different predictions of the information carried by similarity structure of neural signal responses and generated correlational maps (*r*-maps). Finally, these *r*-maps were converted to *z*-maps using Fisher transformation. All the z-maps were then submitted to a group-level one-sample *t*-test to identify voxels in which the similarity between the predicted RDM and observed neural RDM was greater than zero. This allowed us to identify voxels in which information of TD at retrieval might be represented (see **S3 Fig**). The statistical threshold was set as identical to those employed in the univariate analysis, which was at *p* < 0.05 (FWE corrected) at cluster level and *p* < 0.001 at an uncorrected peak level.

#### Leave-one-subject-out approach (LOSO), functional and anatomical ROIs

We applied a LOSO approach to create functional ROIs to avoid statistical bias (52). For instance, in order to identify an ROI (i.e., conjunction mask in **Fig 3D**) for Subj01, we estimated the contrast using a one-sample *t*-test on the whole-brain searchlight *z*-maps obtained from Subj02 to Subj17. Likewise, we also estimated the contrast using a one-sample *t*-test on the contrast maps obtained from the Pmod analysis of Subj02 to Subj17. We set the same threshold reported above to extract clusters from these two statistical maps. We then overlaid the two resultant maps and extracted a conjunction region (mask01), with which we used to extract the value in the searchlight *z*-map from Subj01 for further statistical analysis. This procedure was repeated 17 times and generated 17 different ROIs, which provided statistically independent regions to extract values for testing differences between conditions. For the anatomical ROIs (depicted in **Fig 5** and **S7 Fig**), 7 regions (bilateral precuneus, bilateral angular gyrus, bilateral hippocampus and occipital cortex) of AAL template (53) were created as masks. We extracted and averaged the similarity value within these masks for each subject for statistical tests.

### Candidate representational dissimilarity matrices (RDM)

*Model 1* (TD RDM, 60 × 60). We ranked the difference across the 60 TD levels, from the shortest to the longest TD, for the Within-chapter condition (or Across-chapter condition, the two RDMs were identical because of their matched TD). We first log-transformed the subject-specified TD values for each pair of images and then computed the differences with and among every other TD levels producing 60 × 59/2 values, which were then assigned to the corresponding cells of the RDM.

*Model 2* (Situational Change RDM, 60 × 60). Since the temporal and spatial dimensions were closely inter-correlated. We checked whether the situational change might influence the neural patterns in those voxels that represent the TD information. We analyzed the subject-specific videos frame by frame and marked out the boundaries at which a situational location had changed (see illustration in **S4 Fig**). Then we computed the numbers of situational changes contained in each of the paired images and then computed the differences with and among every other conditions producing 60 × 59/2 values, which were then assigned to the corresponding cells of the RDM.

*Model 3* (RGB-cross-correlation RDM, 60 × 60), *Model 4* (RGB-intensity RDM, 60 × 60) and *Model 5* (RGB-histogram RDM, 60 × 60) considered the perceptual characteristics of the images used in TOJ. For Model 3, the similarity measure was based on the cross-correlation value between two images (image of size 1920 × 1080) for the three color channels (red, green, and blue; RGB). For every pair of images in each of the three color channels (RGB), we computed the cross-correlation coefficients between the pair. This is a measure of the displacement of one image relative to the other; the larger the cross-correlation coefficient (which ranges between −1 and 1), the more similar the two images was. We then computed the differences with and among every other conditions producing 60 × 59/2 values, which were then assigned to the corresponding cells of the RDM. For Model 4, we computed the pixel-wise difference between pair images for the three color channels (RGB). The computed difference is useful when the compared images are taken from a stationary camera with infinitesimal time difference. The output pixel for each color channel is assigned with the value 1 if the absolute difference between the corresponding pixels in the image pair is non-zero, or a value of 0 otherwise. A single value is generated for each of the three color channels by summing all the output pixel values (either 0 or 1). We averaged the sum of difference for all three color-channels for the intensity value of each pair of images and then computed the differences with and among every other conditions producing 60 × 59/2 values, which were then assigned to the corresponding cells of the RDM. For Model 5, we constructed color histograms for image pairs and computed the Sum-of-Square-Difference (SSD) error between them for the three color channels (RGB). For each color channel the intensity values range from 0 to 255 (i.e., 256 bins), we first computed the total number of pixels at each intensity value and then computed the SSD for all 256 bins for each image pair. The smaller the value of the SSD, the more similar the two images (image pair) was. We then computed the differences with and among every other conditions producing 60 × 59/2 values, which were then assigned to the corresponding cells of the RDM. In contrast to model 4, this approach does not require corresponding pixels in the image pair to be the same, but rather measures the existence of pixel intensity in both images. Overall, the three perceptual-similarity models (3, 4 and 5) look at different similarity measures and they complement each other; thus any difference in the appearance of the two images irrespective of the temporal distance, could be accounted for by at least one of the three models. For any two similar images, the RGB-intensity RDM results in a very small value, thus the corresponding pixels are virtually the same for the entire image. The RGB-histogram will also result in a small value as the image pairs will have the same histogram bins. The RGB-cross-correlation value will be close to 1, signifying the similarity in the images. When subsections of a scene are visible in both images with varied brightness, the RGB-cross-correlation value will still be closer to 1 but with a very high RGB-intensity RDM value.

*Model 6* (Scale Invariant Feature Transform (SIFT) RDM, 60 × 60), *Model 7* (Speeded Up Robust Features (SURF) RDM, 60 × 60) and *Model 8* (Histogram of Oriented Gradients (HOG), 60 × 60): For model 6, SIFT transform images into scale-invariant vectors which encoded interest points. For any pair of images to be compared, a computational search over selected scales and image locations is conducted using Difference-of-Gaussian (DoF) to identify potential interest points that are invariant to scale and orientation in each image. Interest points from each image are stabilized using Taylor series expansion of scale space to get a more accurate location of extrema. This is followed by the construction of an orientation histogram to achieve invariance to rotation. Key-points between the two images are then matched by identifying their nearest neighbors using minimum Euclidean distance between the invariant descriptor vectors extracted from each image. For model 7, rather than the use of DoF in SIFT (26) as an approximation of Laplacian-of-Gaussian (LoG), SURF (27) uses Box Filter which is calculated in parallel using integral images (54) to approximate LoG. Wavelet responses in both horizontal and vertical directions are used to assign orientation in SURF. Like SIFT, the first step in SURF consists of fixing a reproducible orientation based on information from a circular region around the interest point (27). A descriptor vector is generated around the interest point using integral image, which is compared to descriptor vectors extracted from a compared image to find a match. The Euclidean distance has been used to measure the similarity between two descriptor vectors from the two images. It is worth noting that SURF was not used in neither Greene et al. (24) nor Aminoff et al (25) but it is robust and much faster to compute than SIFT. However, Greene et al. (24) trained a state-of-the-art convolutional neural network (CNN) on ImageNet 2012 to find visual features to perform their categorization. To remove the time for training a supervised network like CNN which uses SIFT/SURF features in it layers, our dissimilarity measures are based on the features extracted solely from the input images with no prior training. For model 8, in the case of the HOG feature descriptor, we constructed a histogram of directions of gradient over fixed sized grid across the entire image. A vector is generated from each grid cell and correlated with HOG features from another image. It is worth noting that gradients or derivatives of x and y (location) in an image are useful for interest point localization because they have higher values around edges and corners, which hold more information about objects.

*Model 9* (RT RDM, 60 × 60) and *Model 10* (ACC RDM, 60 × 60) were computed based on the behavioral performance of reaction times and percentage correct (0 or 1) for each trial. We averaged the values along the 60 TD levels first (4 repetitions when Within-chapter and Across-chapter conditions collapsed; 2 repetitions for Within-chapter or Across-chapter alone) and computed the differences with and among every other TD levels producing 60 × 59/2 values, which were then assigned to the corresponding cells of the RT RDM and ACC RDM.

## Supporting Information

**S1 Fig.**
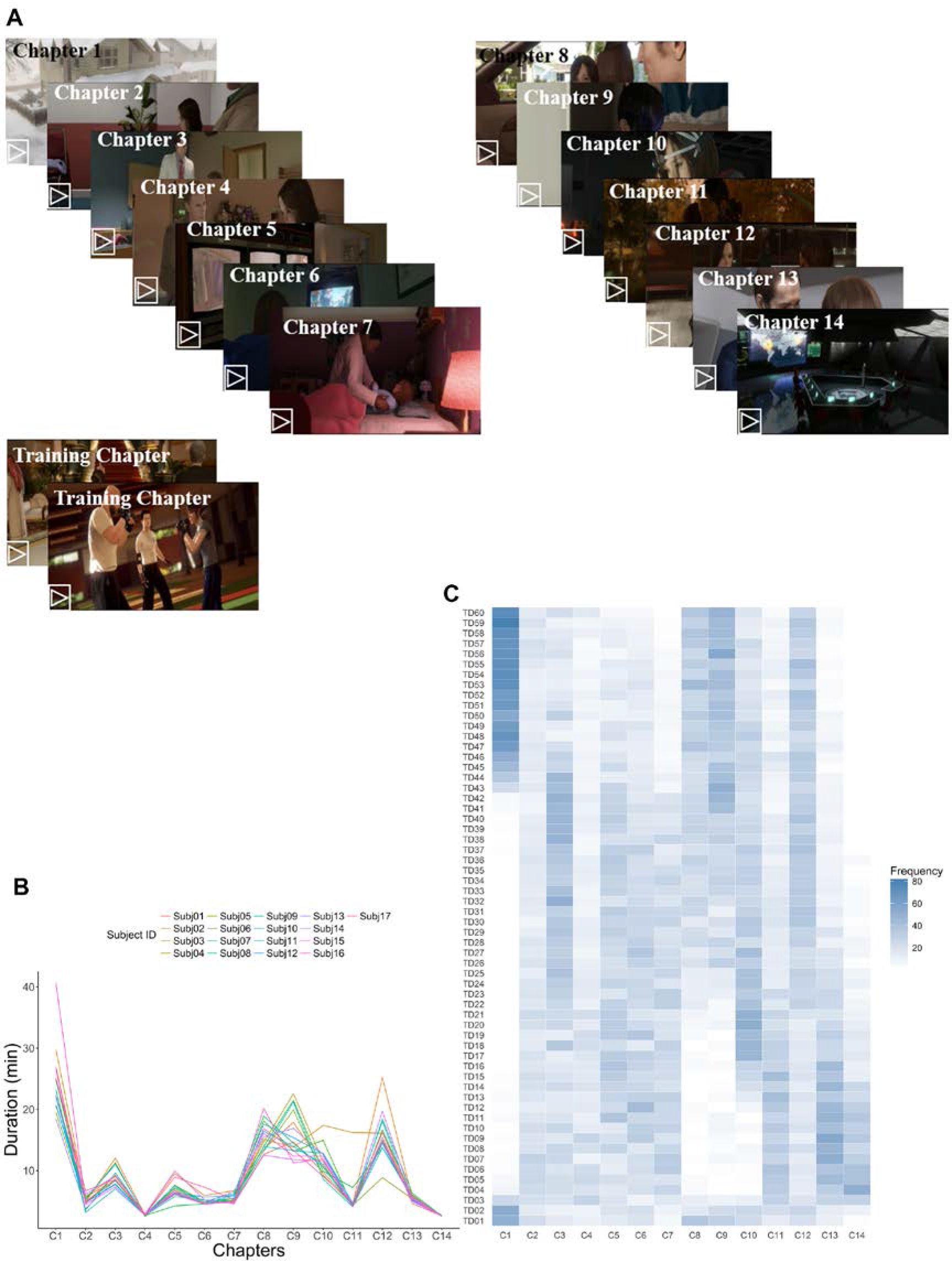
Stimuli. **(A)** 14 different chapters were used: seven in each of two within-subjects TMS sessions (see also **S1 Table**). **(B)** Playing durations varied across chapters and subjects. **(C)** Intensity map showed that the numbers of trials extracted from the 14 game chapters separately for the 60 TD levels across subjects. The numbers of image pairs extracted for the TOJ task were approximately equal across the chapters, except that longer TD are more likely to be extracted from the longest chapter (e.g., C1).

**S2 Fig.**
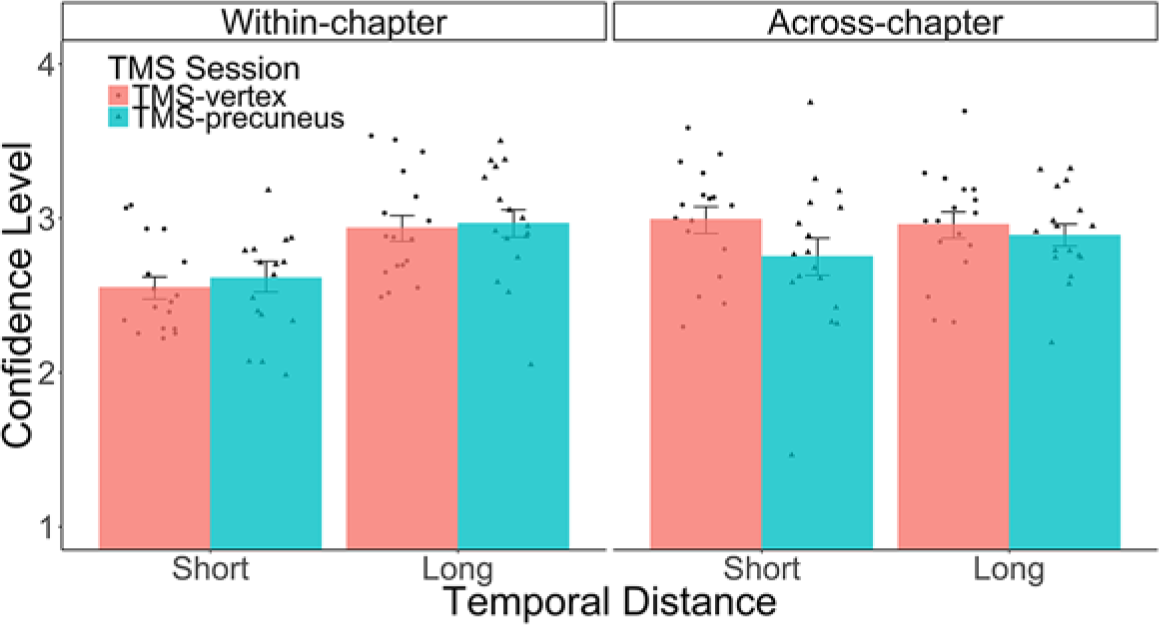
Behavioral results of confidence level. TMS did not result in differences in participants’ confidence level, except for a significant TD × Context interaction (*F*_(1, 16)_ = 19.12, *P* < 0.001, *η*^2=^ 4.52%), which was driven by significant differences in confidence level between short and long TD in Within-chapter condition (*t*_(33)_ = 6.76, *P* < 0.001), but not in Across-chapter condition (*t*_(33)_ = 1.17, *P* = 0.250). Dots indicate behavioral performance per subject. Error bars denote the SEM over subjects.

**S3 Fig.**
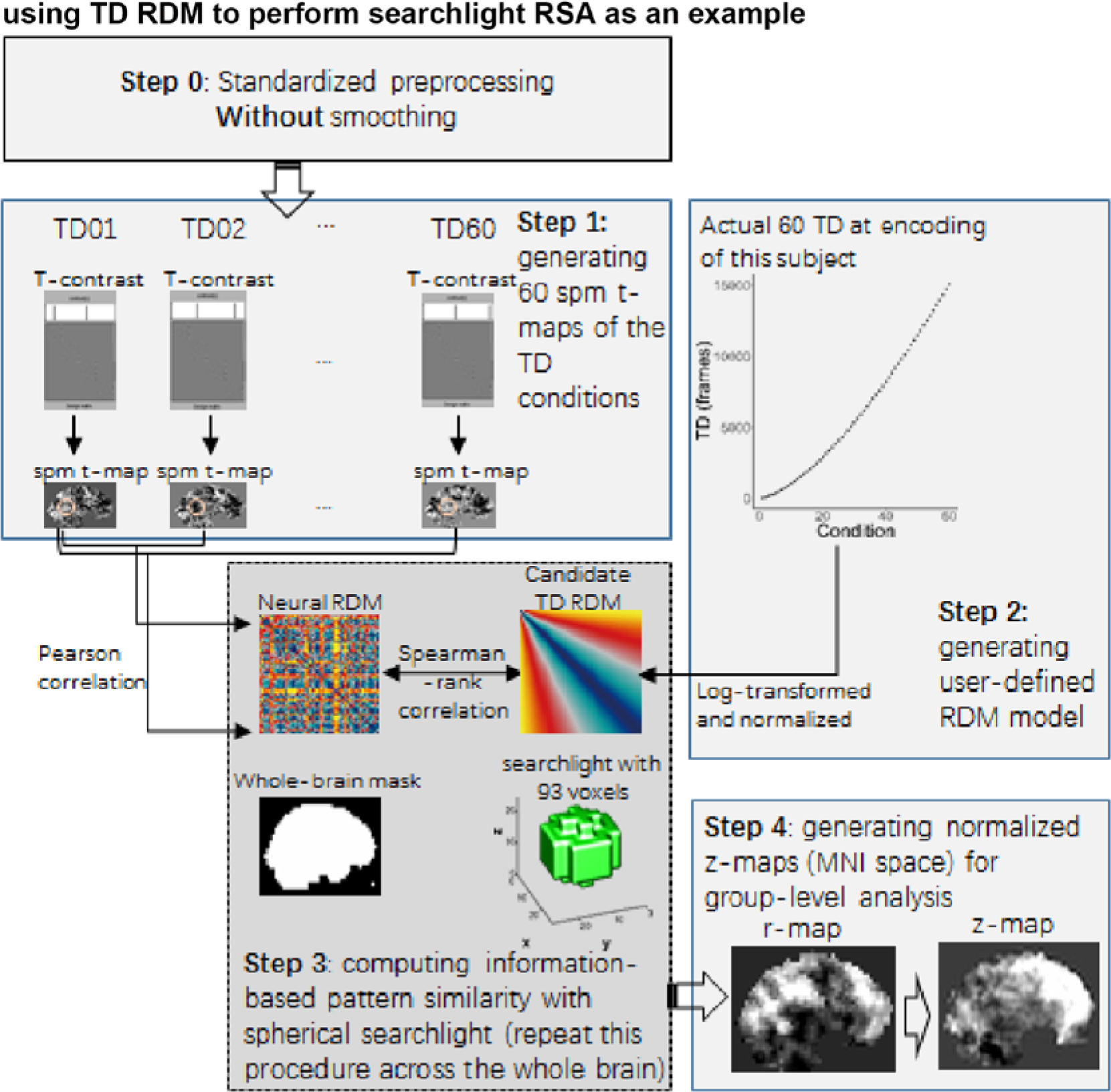
Pipeline of RSA analysis. Using one model (TD RDM) to perform the searchlight procedure as an example in one subject (Subj01). The same principle and procedure applied to all other candidate RDMs and subjects.

**S4 Fig.**
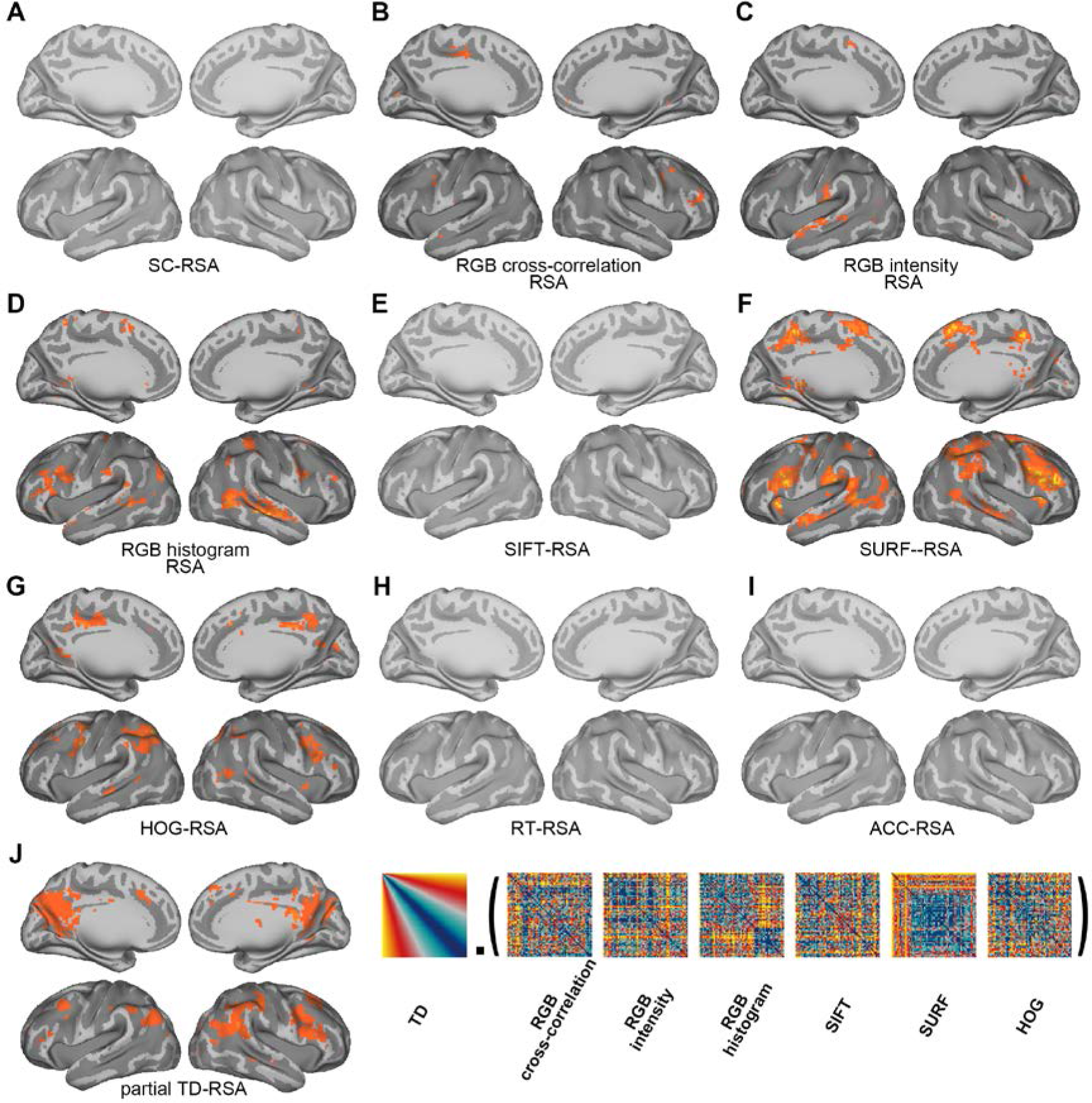
RSA searchlight results for TMS-vertex sessions using candidate RDMs. **(A)** Situational Change RDM (see **S5 Fig**). **(B)** RGB cross-correlation RDM. **(C)** RGB-intensity RDM. **(D)** RGB-histogram RDM. **(E)** SIFT RDM. **(F)** SURF RDM. **(G)** HOG RDM. **(H)** Reaction time RDM. **(I)** Accuracy RDM. **(J)** A partial correlation analysis showed that the TD representation in the posteromedial parietal areas was not subject to influences due to the six perceptual RDMs. MRI results are displayed at *P*_uncorrected_ < 0.001.

**S5 Fig.**
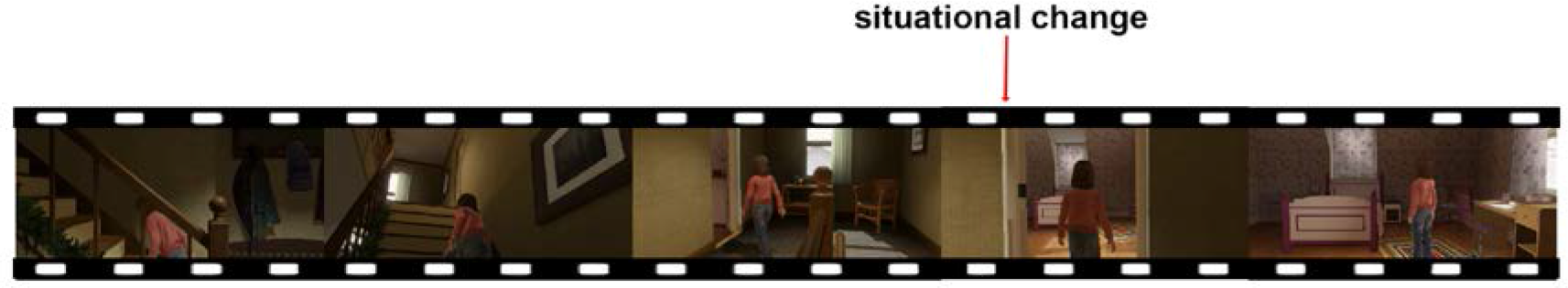
An excerpt of a subject-specific video. An excerpt of a subject-specific video illustrates how situational changes were defined frame-by-frame for Situational Change RDMs and pmod analyses. Noting that time and space in our task were partially correlated, we quantified subject-specific situational changes in terms of the number of locations each participant had traversed in the video (i.e., spatial displacement embedded in the paired images presented in TOJ task).

**S6 Fig.**
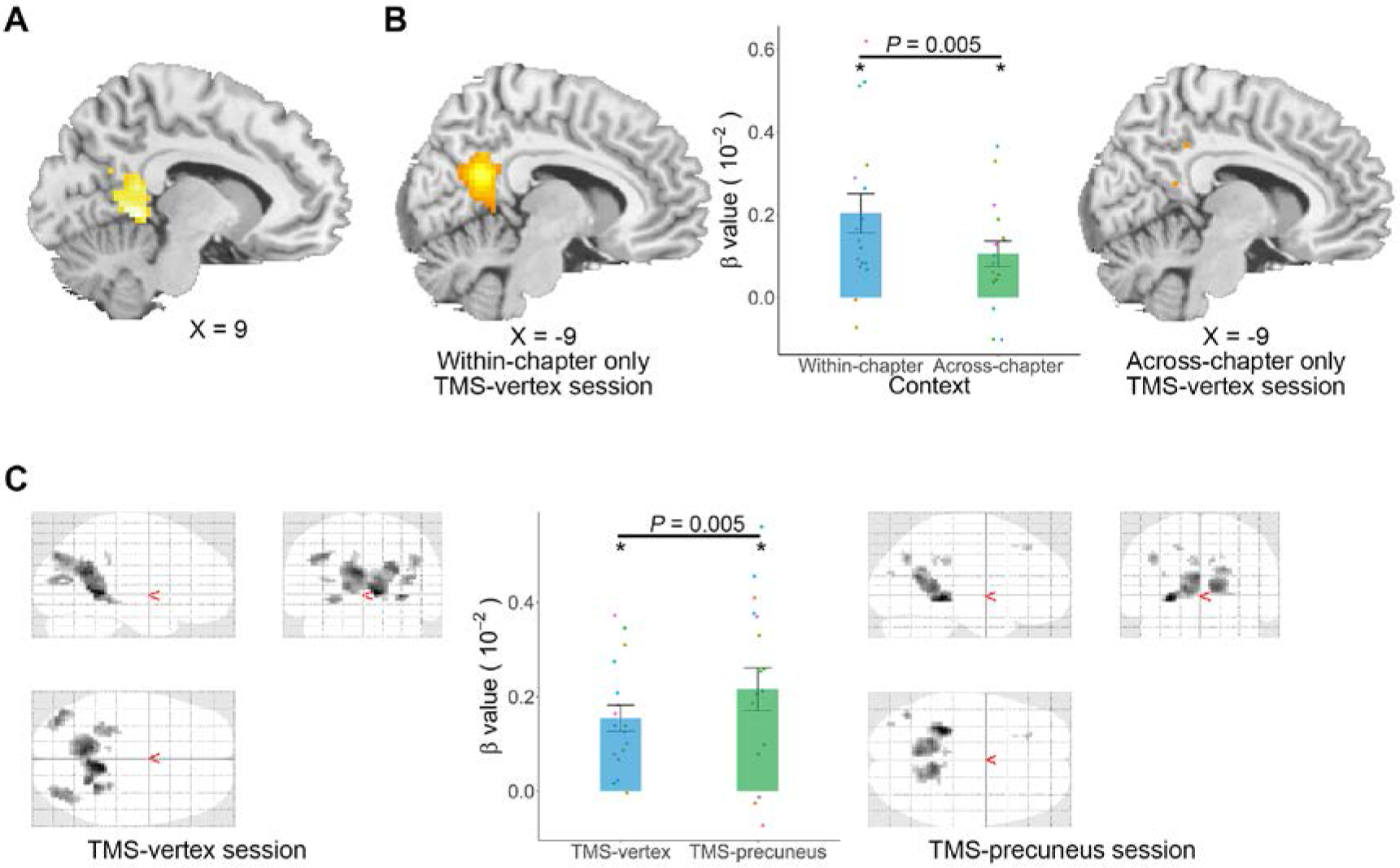
Pmod analyses for TMS-vertex session and TMS-precuneus session. **(A)** Temporal distance (TD) and reaction time (RT) as modulatory regressors for TMS-vertex session (Within-chapter and Across-chapter trials collapsed). Having removed the influence of RT, neural activity in the precuneus/retrosplenial cortex and posterior hippocampus still increased with TD. **(B)** TD and RT as modulatory regressors separately for Within-chapter (left) and Across-chapter (right) conditions in the TMS-vertex session. After having removed the influence of RT, neural activity in the precuneus, extending into the retrosplenial cortex and posterior cingulate cortex, remained linearly associated with TD in Within-chapter condition. The *β* values extracted from the cluster was also higher in the Within-chapter condition than in the Across-chapter condition (*P* = 0.005). **(C)** In contrast to the removal of the multivoxel representation, TMS to the precuneus did not impact on the activation intensity in the precuneus and retrosplenial cortex (right). Moreover, activation in these clusters (containing the precuneus) increased with temporal distance irrespective of TMS Precuneus and temporal context memory stimulation (middle panel). Activation maps are displayed at *P*_uncorrected_ < 0.001.

**S7 Fig.**
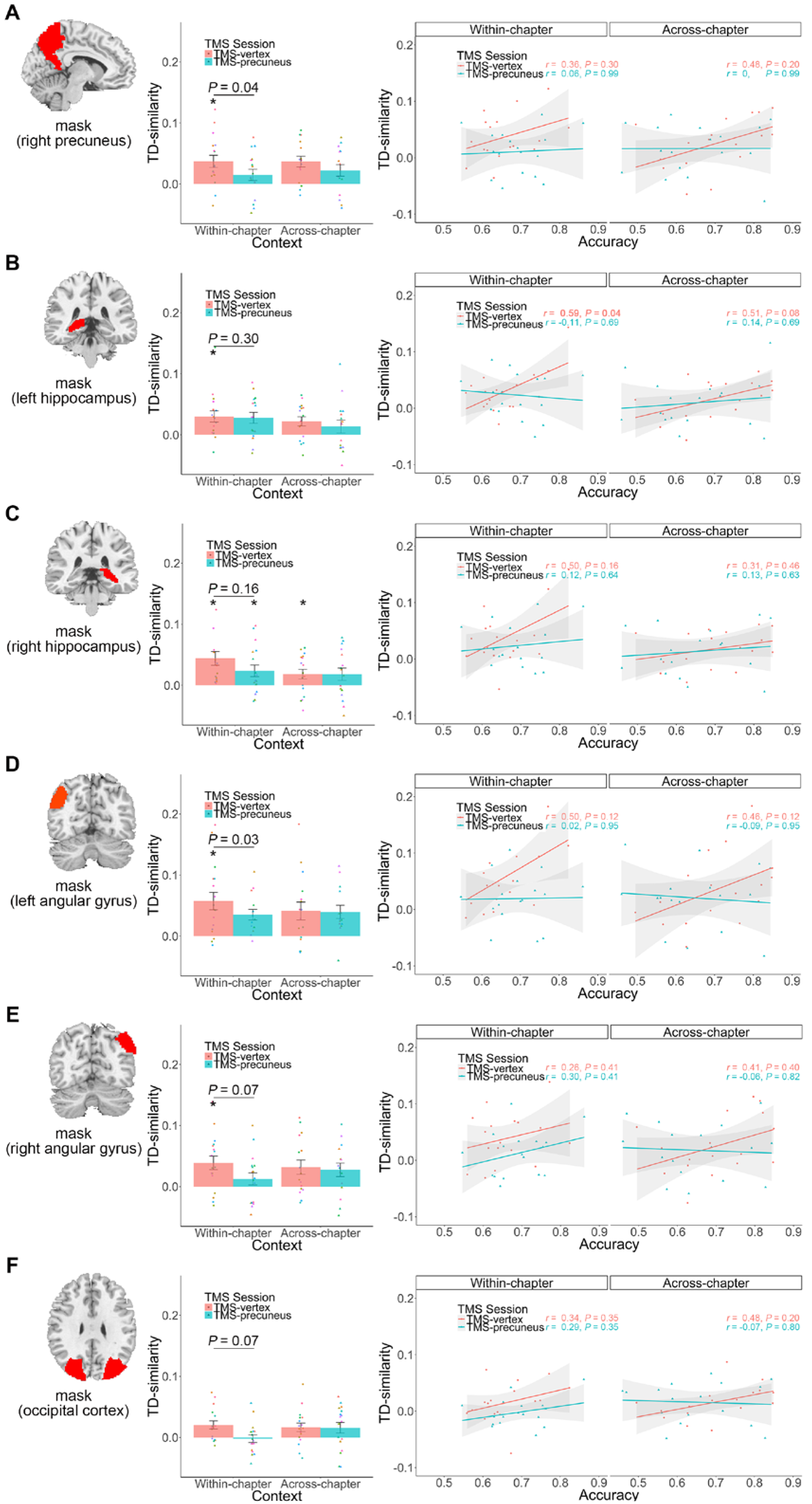
TD-similarity indices in six anatomical ROIs. (**A**: right precuneus, **B-C**: bilateral hippocampus, **D-E**: bilateral angular gyrus, and **F**: whole occipital cortex) separately for Within-chapter and Across-chapter conditions, in both TMS sessions, and their correlation with individual subjects TOJ accuracy. * *P* < 0.05, FDR-corrected, one-sample *t*-test against zero. Horizontal lines indicate paired *t*-test between two bars and *P* values are FDR-corrected.

**S8 Fig.**
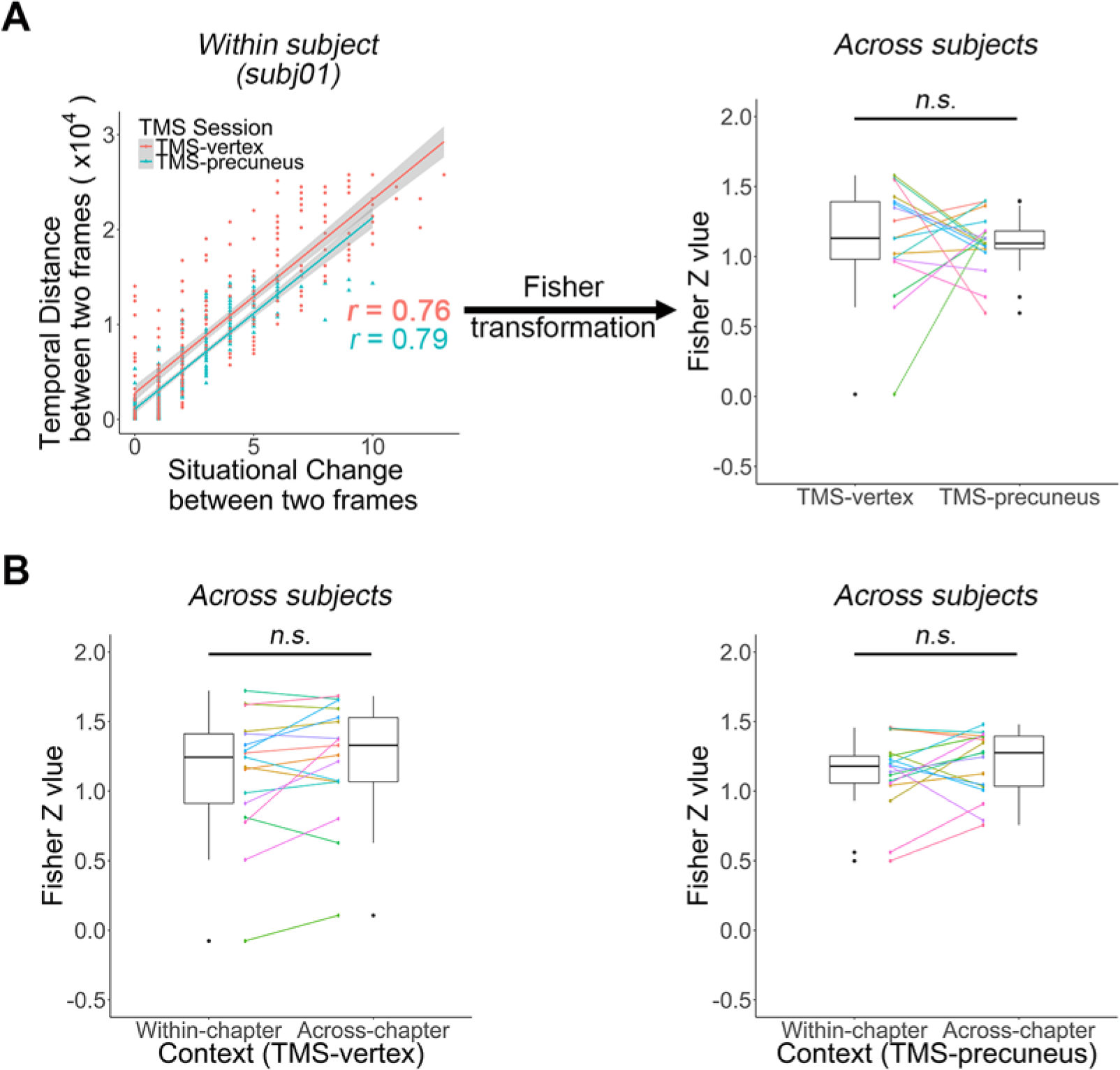
Correlation coefficients. Correlation coefficients were very high between temporal distance and the number of situational changes across trials in all subjects. **(A)** Correlation coefficients are highly correlated for all participants in both TMS sessions (each dot represents a trial; one participant illustrated as an example, left panel). Fisher *z* values (transformed from correlation coefficients) for all participants are shown in a boxplot on the right side. Fisher z values of two conditions are significantly greater than zero (TMS-vertex session: mean ± se = 1.12 ± 0.1, *t*_16_ = 11.52, *P* = 3.71 ×10^−9^; TMS-precuneus session: mean ± se = 1.10 ± 0.05, *t*_16_ = 21.08, *P* = 4.24 × 10^−13^). No significant differences exist between the two sessions (*P* = 0.83). Individual participants’ values are connected between the two boxplots. (B) For completeness, we also calculated the Fisher *z* values for all participants in the Across-chapter and Within-chapter conditions of two TMS sessions separately. *Z* values are significantly greater than zero in each conditions (*P* < 0.01). No significant differences exist between the two sessions (*P* > 0.05). n.s. denotes not significant.(TIF)

**S1 Table.** Descriptive statistics of TMS motor threshold and full counter-balancing for gameplay chapters by TMS sites.

| ID | Motor<br>thresh<br>old | Session 1 |  |  | Delay<br>(days) | Session 2 |  |  |
| --- | --- | --- | --- | --- | --- | --- | --- | --- |
|  |  | Chapters<br>(Playing) | Duration<br>(min) | TMS Site |  | Chapters<br>(Playing) | Duration<br>(min) | TMS Site |
| Subj01 | 66% | Chap 1 ~ 7 | 89.39 | Precuneus | 9 | Chap 8 ~ 14 | 35.19 | Vertex |
| Subj02 | 67% | Chap 1 ~ 7 | 102.70 | Vertex | 6 | Chap 8 ~ 14 | 44.46 | Precuneus |
| Subj03 | 80% | Chap 8 ~ 14 | 35.53 | Vertex | 9 | Chap 1 ~ 7 | 114.21 | Precuneus |
| Subj04 | 58% | Chap 8 ~ 14 | 36.41 | Vertex | 2 | Chap 1 ~ 7 | 94.70 | Precuneus |
| Subj05 | 65% | Chap 1 ~ 7 | 80.50 | Precuneus | 8 | Chap 8 ~ 14 | 35.31 | Vertex |
| Subj06 | 67% | Chap 1 ~ 7 | 95.05 | Vertex | 3 | Chap 8 ~ 14 | 39.63 | Precuneus |
| Subj07 | 72% | Chap 8 ~ 14 | 32.41 | Vertex | 4 | Chap 1 ~ 7 | 97.06 | Precuneus |
| Subj08 | 73% | Chap 1 ~ 7 | 96.08 | Precuneus | 8 | Chap 8 ~ 14 | 37.03 | Vertex |
| Subj09 | 69% | Chap 1 ~ 7 | 88.81 | Vertex | 7 | Chap 8 ~ 14 | 31.46 | Precuneus |
| Subj10 | 80% | Chap 8 ~ 14 | 35.69 | Precuneus | 7 | Chap 1 ~ 7 | 89.79 | Vertex |
| Subj11 | 65% | Chap 8 ~ 14 | 38.77 | Vertex | 26 | Chap 1 ~ 7 | 86.02 | Precuneus |
| Subj12 | 71% | Chap 1 ~ 7 | 90.56 | Precuneus | 16 | Chap 8 ~ 14 | 35.89 | Vertex |
| Subj13 | 68% | Chap 8 ~ 14 | 35.66 | Vertex | 7 | Chap 1 ~ 7 | 87.98 | Precuneus |
| Subj14 | 73% | Chap 1 ~ 7 | 87.94 | Precuneus | 7 | Chap 8 ~ 14 | 40.56 | Vertex |
| Subj15 | 57% | Chap 8 ~ 14 | 32.72 | Precuneus | 5 | Chap 1 ~ 7 | 91.46 | Vertex |
| Subj16 | 69% | Chap 8 ~ 14 | 35.44 | Vertex | 5 | Chap 1 ~ 7 | 108.49 | Precuneus |
| Subj17 | 61% | Chap 1 ~ 7 | 91.10 | Precuneus | 6 | Chap 8 ~ 14 | 42.13 | Vertex |

**S2 Table.** Brain representation associated with TD RDM in TMS-vertex session.

| Cluster |  | peak |  |  | % Cluster | Brain regions |
| --- | --- | --- | --- | --- | --- | --- |
| k | p-FWE | X | y | z |  |  |
| 7554 | <.001 | 30 | 21 | 38 | 5.41 | Precuneus_L |
|  |  |  |  |  | 5.32 | Precuneus_R |
|  |  |  |  |  | 4.38 | Frontal_Mid_2_R |
|  |  |  |  |  | 3.34 | Angular_R |
|  |  |  |  |  | 3.24 | Parietal_Inf_L |
|  |  |  |  |  | 3.15 | Frontal_Inf_Tri_R |
|  |  |  |  |  | 2.99 | Parietal_Inf_R |
|  |  |  |  |  | 2.91 | Frontal_Mid_2_L |
|  |  |  |  |  | 2.83 | Frontal_Inf_Tri_L |
|  |  |  |  |  | 2.74 | Angular_L |
|  |  |  |  |  | 2.73 | SupraMarginal_R |
|  |  |  |  |  | 2.73 | Frontal_Inf_Oper_R |
|  |  |  |  |  | 2.65 | Frontal_Sup_2_R |
|  |  |  |  |  | 2.61 | Cuneus_L |
|  |  |  |  |  | 2.58 | Cingulate_Mid_R |
|  |  |  |  |  | 2.57 | Postcentral_R |
|  |  |  |  |  | 2.55 | Temporal_Mid_R |
|  |  |  |  |  | 2.26 | Calcarine_L |
|  |  |  |  |  | 2.22 | Cuneus_R |
|  |  |  |  |  | 1.87 | Precentral_R |
|  |  |  |  |  | 1.85 | Occipital_Mid_R |
|  |  |  |  |  | 1.62 | Calcarine_R |
|  |  |  |  |  | 1.62 | Frontal_Sup_Medial_R |
|  |  |  |  |  | 1.56 | Cingulate_Mid_L |
|  |  |  |  |  | 1.52 | Occipital_Sup_R |
|  |  |  |  |  | 1.47 | Frontal_Sup_Medial_L |
|  |  |  |  |  | 1.42 | Temporal_Sup_R |
|  |  |  |  |  | 1.22 | Occipital_Mid_L |
|  |  |  |  |  | 1.02 | Cingulate_Post_L |
|  |  |  |  |  | 1.01 | Precentral_L |

Note. % Cluster denotes percentage of voxels of the cluster that is contained in each of the anatomical regions (53).

**S3 Table.** Brain activation parametrically modulated by temporal distance (TD) in TMS-vertex session (see Fig 3C)

| Regressor | Cluster |  | Voxel |  |  |  | Brain Regions |
| --- | --- | --- | --- | --- | --- | --- | --- |
|  | k | p-FWE | Z value | x | Y | z |  |
| TD | 555 | <.001 | 4.62 | 9 | -45 | -1 | Cerebelum_4_5_R |
|  |  |  | 4.35 | -6 | -57 | 20 | Precuneus_L |
|  |  |  | 4.27 | 6 | -54 | 5 | Vermis_4_5 |
|  | 116 | .002 | 3.87 | 36 | -81 | 14 | Occipital_Mid_L |
|  |  |  | 3.78 | 42 | -72 | 29 | Occipital_Mid_L |
|  |  |  | 3.75 | 36 | -69 | 11 | Occipital_Mid_R |
|  | 117<sup>†</sup> | .01 | 4.34 | 45 | -36 | 5 | Temporal_Sup_R |
|  |  |  | 4.17 | 54 | -36 | 5 | Temporal_Sup_R |
|  |  |  | 3.81 | 48 | -27 | -1 | Temporal_Sup_R |

Note. <sup>†</sup> denotes a negative relationship between TD and brain activity.

### S1 Video. An excerpt of the gameplay video

(MP4)

## Acknowledgements

This research is sponsored by the National Natural Science Foundation of China 31872783 (Y.H.), by the Ministry of Education of PRC Humanities and Social Sciences Research grant 16YJC190006, STCSM Shanghai Pujiang Program 16PJ1402800, STCSM Natural Science Foundation of Shanghai 16ZR1410200, Large Instruments Open Foundation (ECNU), and NYU Shanghai and the NYU-ECNU Institute of Brain and Cognitive Science at NYU Shanghai (S.C.K.).

## Author contributions

Q.Y. designed and conducted the study, analyzed data, drafted and wrote the manuscript. Y.H. discussed the results and commented on drafts. Y.K. advised on TMS protocol. K.A. produced indices for RDM Models 4, 5, 6, 7 and 8. S.C.K. designed the study, supervised the research, and wrote the manuscript.

## Competing interests

The authors declare no competing interests.

## Code availability

The codes used for the analyses are available upon request.

## Data availability

The data reported in this study are available on http://datadryad.org/review?doi=doi:10.5061/dryad.pj038.

